# No Reliable Evidence Supports the Presence of Javan Tigers - Data Issues Related to the DNA Analysis of a Recent Hair Sample

**DOI:** 10.1101/2024.04.08.588384

**Authors:** Zheng-Yan Sui, Nobuyuki Yamaguchi, Yue-Chen Liu, Hao-Ran Xue, Xin Sun, Philip Nyhus, Shu-Jin Luo

**Affiliations:** The State Key Laboratory of Protein and Plant Gene Research, School of Life Sciences; Peking-Tsinghua Center for Life Sciences, Academy for Advanced Interdisciplinary Studies; Institute of Ecology, Peking University, Beijing 100871, China; Institute of Tropical Biodiversity and Sustainable Development, University of Malaysia Terengganu, Kuala Nerus 21030, Terengganu, Malaysia; Wildlife Conservation Research Unit, Department of Biology, Oxford University, Recanati-Kaplan Centre, Tubney House, Abingdon Road, Tubney, UK. OX13 5QL; Department of Human Evolutionary Biology, Harvard University, Cambridge, MA, 02138, USA; Department of Genetics, Harvard Medical School, Boston, MA, 02115, USA; University of Potsdam, Institute for Biochemistry and Biology, Karl-Liebknecht-Str. 24-25, D-14476 Potsdam-Golm, Germany; Center for Evolutionary Hologenomics, Globe Institute, Faculty of Health and Medical Sciences, University of Copenhagen, DK-1353 Copenhagen, Denmark; Environmental Studies Department, Colby College, Waterville, Maine, 04901 USA

**Keywords:** Javan tiger, Numt, *Panthera tigris sondaica*

## Abstract

A paper recently published in *Oryx* by Wirdateti et al. (2024) suggests that the extinct Javan tiger may still survive on the Island of Java, Indonesia, based on mtDNA analysis of a single hair collected from a claimed tiger encounter site. After carefully re-analyzing the data presented in Wirdateti et al. (2024), we conclude that there is little support for the authors’ statements. Importantly, the sequences of the putative tiger hair and museum Javan tiger specimens generated by the authors are not from tiger cytoplasmic mitochondrial DNA but more likely the nuclear copies of mitochondrial DNA. In addition, the high mismatches found between the two “Javan tiger” sequences generated by the authors is unusual for homologous sequences that are both from tigers and hence indicative of data unreliability. Yet, too few details regarding the quality control were provided in Wirdateti et al. (2024) to rule out the possibility of contamination introduced during the data production process. In conclusion, it is inappropriate to use these unreliable sequences presented in Wirdateti et al. (2024) to infer the existence of the Javan tiger.

---

We are writing with regard of a recent paper published in the Oryx entitled “Is the Javan tiger *Panthera tigris sondaica* extant? DNA analysis of a recent hair sample” by Wirdateti and others (Wirdateti et al., 2024) (https://doi.org/10.1017/S0030605323001400).

We read the article with great excitement, which was, quickly and sadly, replaced by concerns about the credibility of data and hence the reliability of the conclusion. We have three major concerns, which are 1) the sequences that the authors obtained are not genuine tiger mitochondrial DNA (mtDNA), 2) they are likely nuclear pseudogene copies of mitochondrial DNA (Numt), and 3) readers cannot evaluate the original data reliability because few details concerning the quality control are provided in the article. We briefly explain these below.

There has been no confirmed sighting of the Javan tiger since the 1970s (Seidensticker, 1987), and the subspecies was officially categorized as extinct in 2008 by the IUCN Red List of Threatened Species in 2008 (Jackson & Nowell, 2008). Therefore, it was a great surprise to read that a putative encounter with a tiger occurred in 2019 at a community plantation in West Java, and that a single hair sample was collected from a fence nearby, which was then analyzed by the authors. The authors amplified and sequenced a 1,043 bp cytochrome b mtDNA segment from the hair collected, and compared it to those of leopards and origin-known tiger subspecies, including a museum Javan tiger specimen collected in 1930 that the authors amplified and sequenced. Phylogenetic trees showed that the hair sample aligned most closely with the museum Javan tiger specimen, forming a clade distinct from other tiger subspecies and the Javan leopard. Based on the results the authors conclude that the hair belongs to the Javan tiger, implying tigers still survive on the island.

This would be extremely exciting news if genuine. However, very disappointedly, after carefully re-analyzing the data presented by Wirdateti et al. (2024) we conclude that there is no support for the authors’ conclusions because of the following three main reasons.

## 1. The sequences that the authors obtained are not tiger mitochondrial DNA segments

In Wirdateti et al. (2024), the genetic clade including the hair sample in question (NCBI Accession OQ601561.1) and the museum Javan tiger specimen from 1930 (OQ601562.1) is an outgroup to the tiger mtDNA clade and phylogenetically equidistant from both tigers and leopards, which is a pattern that is not observed from the previous studies involving the Javan tiger (Xue et al., 2015; Sun et al., 2023). To investigate this issue, we conducted phylogenetic analysis on the two putative Javan tiger sequences produced by Wirdateti et al. (2024), along with other published mtDNA sequences from *Panthera* species (28 *P. tigris*, three *P. pardus*, three *P. leo*, three *P. onca*, and three *P. uncia*) (Table 1). MUSCLE (v 5.1) was implemented for multi-sequence alignment of the 42 sequences. The alignment was manually trimmed 72 bp from both sides, resulting in a 971 bp nucleotide sequence matrix without gaps of missing data.

**TABLE 1.** Mitogenome DNA sequences involved in this study.

| Sequence ID | Taxonomy | Source |
| --- | --- | --- |
| **Sequences from Wirdateti et al. (2024, N=7)** |  |  |
| OQ601561.1 | *P. tigris sondaica* (putative) | NCBI Accession: OQ601561.1 |
| OQ601562.1 | *P. tigris sondaica* (putative) | NCBI Accession: OQ601562.1 |
| OQ629467.1 | *P. tigris sumatrae* | NCBI Accession: OQ629467.1 |
| OQ629468.1 | *P. tigris sumatrae* | NCBI Accession: OQ629468.1 |
| OQ629469.1 | *P. tigris sumatrae* | NCBI Accession: OQ629469.1 |
| OQ629470.1 | *P. tigris sumatrae* | NCBI Accession: OQ629470.1 |
| OQ629471.1 | *P. tigris sumatrae* | NCBI Accession: OQ629471.1 |
| **Sequences from other tigers (N=24)** |  |  |
| NC_010642.1 | *P. tigris* | NCBI Accession: NC_010642.1 |
| pti183 | *P. tigris sumatrae* | Sun et al. (2023) |
| pti184 | *P. tigris sumatrae* | Sun et al. (2023) |
| pti096 | *P. tigris sumatrae* | Sun et al. (2023) |
| pti105 | *P. tigris tigris* | Sun et al. (2023) |
| pti103 | *P. tigris tigris* | Sun et al. (2023) |
| pti331 | *P. tigris tigris* | Sun et al. (2023) |
| PTV02 | *P. tigris virgata* | Sun et al. (2023) |
| PTV17 | *P. tigris virgata* | Sun et al. (2023) |
| pti305 | *P. tigris corbetti* | Sun et al. (2023) |
| pti306 | *P. tigris corbetti* | Sun et al. (2023) |
| pti307 | *P. tigris corbetti* | Sun et al. (2023) |
| pti247 | *P. tigris jacksoni* | Sun et al. (2023) |
| pti269 | *P. tigris jacksoni* | Sun et al. (2023) |
| pti272 | *P. tigris jacksoni* | Sun et al. (2023) |
| RUSA06_cap | *P. tigris* | Sun et al. (2023) |
| RUSA23_cap | *P. tigris* | Sun et al. (2023) |
| RFET0002 | *P. tigris altaica* | Sun et al. (2023) |
| RFET0007 | *P. tigris altaica* | Sun et al. (2023) |
| pti220 | *P. tigris amoyensis* | Sun et al. (2023) |
| HPS | *P. tigris amoyensis* | Sun et al. (2023) |
| M2 | *P. tigris amoyensis* | Sun et al. (2023) |
| Maza0008 | *P. tigris sondaica* | Sun et al. (2023) |
| Nobb0004 | *P. tigris balica* | Sun et al. (2023) |
| **Sequences from other *Panthera* animals (N=12)** |  |  |
| JF720183.1 | *P. pardus* | NCBI Accession: JF720183.1 |
| MH588626.1 | *P. pardus* | NCBI Accession: MH588626.1 |
| NC_010641.1 | *P. pardus* | NCBI Accession: NC_010641.1 |
| NC_028302.1 | *P. leo* | NCBI Accession: NC_028302.1 |
| KP001504.1 | *P. leo* | NCBI Accession: KP001504.1 |
| KP001505.1 | *P. leo* | NCBI Accession: KP001505.1 |
| NC_022842.1 | *P. onca* | NCBI Accession: NC_022842.1 |
| KM236783.1 | *P. onca* | NCBI Accession: KM236783.1 |
| KF483864.1 | *P. onca* | NCBI Accession: KF483864.1 |
| NC_010638.1 | *P. uncia* | NCBI Accession: NC_010638.1 |
| MT423723.1 | *P. uncia* | NCBI Accession: MT423723.1 |
| MT423722.1 | *P. uncia* | NCBI Accession: MT423722.1 |

A maximum likelihood phylogenetic tree was constructed using IQ-TREE (v 2.3.0), with the HKY+G model selected by jModelTest (v 2.1.10) and statistical support was evaluated based on 10,000 bootstraps. *Prima facie*, our results (Fig. 1) appear to recapitulate the pattern documented by Wirdateti et al. (2024), in which the clade including OQ601561.1 and OQ601562.1 is an outgroup of the tiger mtDNA clade. However, the clade exhibits an unusually elongated branch length in comparison to those amongst all other tiger subspecies. This pattern is not observed in previous studies based on partial (Xue et al., 2015) or full (Sun et al., 2023) mtDNA sequences from origin-known Javan tiger specimens, and therefore, strongly suggests that the two sequences generated by the authors do not originate from Javan tiger mtDNA.

**FIG. 1.**
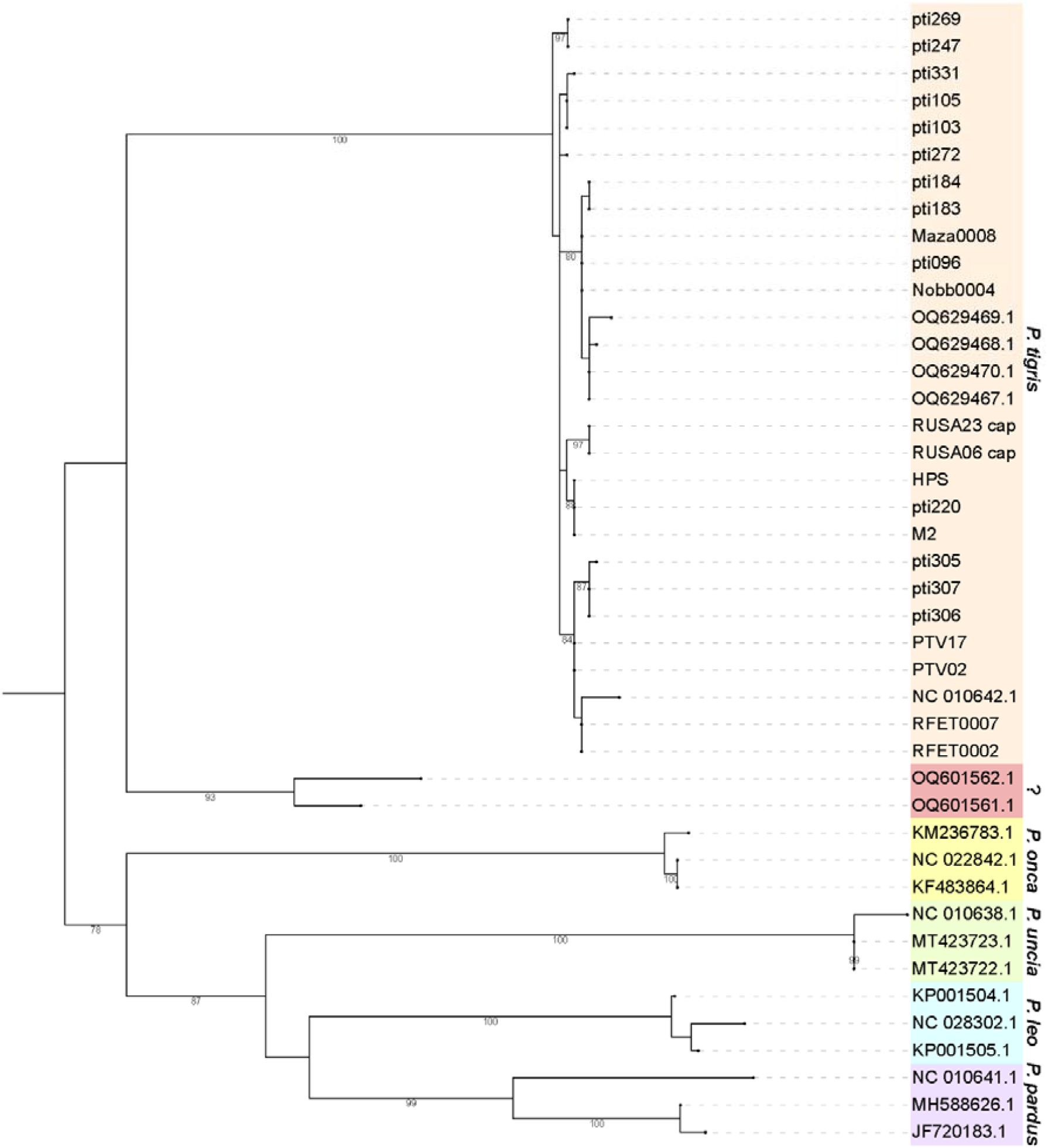
ML phylogeny inferred from the 971 bp region in mitochondria cytochrome b sequence. Sequences are marked with different colors based on their taxonomic classification, and the species are labeled alongside. The suspicious sequences (OQ601561.1 and OQ601562.1) are annotated with a question mark. The support rates over 70% in 10,000 bootstrap tests were labeled near the nodes. The *P. tigris* sequences included are from 9 different subspecies, as listed in Table 1.

We further evaluated the pair-wise genetic distances (p-distance) among the sequences using Biopython (v1.83). The average distance among the 28 published tiger mtDNA sequences is 5.645×10^-3^ (378 pairwise distance calculations, SD = 2.733×10^-3^), while the average distance between the “Javan tiger” sequences generated by the authors and the published tiger mtDNA sequences is 0.07353 (56 pairwise distance calculations, SD = 2.872×10^-3^), which is 13 times as large as the average between-tiger genetic distance. For comparison, the average mtDNA genetic distance between a non-tiger *Panthera* species and a tiger is 0.1049 (336 pairwise distance calculations, SD = 4.854×10^-3^), only slightly larger than the level of genetic distance between the “Javan tigers” and published tiger subspecies.

From both the perspective of phylogenetic pattern and genetic distance, the two “Javan tiger” sequences generated by the authors exhibit significant disparities from the mtDNA sequences of all tiger subspecies, including the published Javan tiger mtDNA haplotype. Such differences compel us to cast doubts on the genuine mtDNA origin of the two “Javan tiger” sequences. It is highly improbable for these two sequences to originate from tiger mtDNA (let alone Javan tiger). That would explain why the two sequences do not cluster with other tigers.

## 2. The sequences that the authors obtained are likely nuclear mitochondrial DNA segments

Nuclear mitochondrial DNA pseudogene segments (Numts) are results of transfer of cytoplasmic mtDNA (Cymt) copies into the nuclear DNA, a common scenario found in tiger and *Panthera* species genomes (Luo et al., 2004; Kim et al., 2006). Given their common origin, there is the possibility that both Cymt and Numt segments are amplified by non-specific primers.

A BLASTn (v2.14.1) search against the latest tiger genome assembly pti1_mat1.1 (NCBI Accession GCF_018350195.1) indicates the Numt as the most likely sources of OQ601561.1 and OQ601562.1. The best- (NC_056676.1:5,568,675-5,569,645) and second-best (NC_056676.1:5,585,283-5,586,253) matched regions of OQ601561.1 and OQ601562.1 are both located on an autosomal scaffold corresponding to tiger chromosome F2. The nucleotide sequence identities of these matches are all over 97.4% across the 971 bp trimmed sequences, whereas the similarity to the tiger mtDNA (NC_010642.1) is 92.5% or below. By contrast, mtDNA segments from a previously published Javan tiger (Maza0008) (Sun et al., 2023), and the Sumatran tigers acquired by the authors matched the tiger mtDNA, with over 98.75% sequence similarity (Fig. 1, Table 2). These results suggested that the two “Javan tiger” sequences generated by the authors are not derived from the Cymt, but more likely, the Numt.

**TABLE 2.** BLAST results on the tiger reference genome.

| Query | Subject | % identity | Align-<br>ment<br>length | Mis-<br>matches | Gaps | Start in<br>query | End in<br>query | Start in<br>subject | End in<br>subject | E-value | Bit<br>score |
| --- | --- | --- | --- | --- | --- | --- | --- | --- | --- | --- | --- |
| OQ601561.1 | NC_056676.1 | 97.528 | 971 | 24 | 0 | 1 | 971 | 5568675 | 5569645 | 0 | 1661 |
| OQ601561.1 | NC_056676.1 | 97.425 | 971 | 25 | 0 | 1 | 971 | 5585283 | 5586253 | 0 | 1655 |
| OQ601561.1 | NC_010642.1 | 92.482 | 971 | 73 | 0 | 1 | 971 | 15188 | 16158 | 0 | 1389 |
| OQ601561.1 | NC_056668.1 | 87.449 | 972 | 120 | 2 | 1 | 971 | 24377331 | 24378301 | 0 | 1118 |
| OQ601561.1 | NC_056677.1 | 82.851 | 968 | 159 | 4 | 1 | 968 | 68227587 | 68228547 | 0 | 861 |
| OQ601561.1 | NC_056663.1 | 85.845 | 763 | 89 | 4 | 1 | 763 | 107153940 | 107154683 | 0 | 793 |
| OQ601561.1 | NC_056660.1 | 79.915 | 946 | 166 | 18 | 33 | 971 | 897423 | 898351 | 0 | 673 |
| OQ601561.1 | NC_056666.1 | 79.239 | 973 | 190 | 7 | 1 | 971 | 58091713 | 58092675 | 0 | 667 |
| OQ601562.1 | NC_056676.1 | 97.528 | 971 | 24 | 0 | 1 | 971 | 5568675 | 5569645 | 0 | 1661 |
| OQ601562.1 | NC_056676.1 | 97.425 | 971 | 25 | 0 | 1 | 971 | 5585283 | 5586253 | 0 | 1655 |
| OQ601562.1 | NC_010642.1 | 92.181 | 972 | 74 | 2 | 1 | 971 | 15188 | 16158 | 0 | 1373 |
| OQ601562.1 | NC_056668.1 | 87.879 | 957 | 113 | 3 | 1 | 955 | 24377331 | 24378286 | 0 | 1122 |
| OQ601562.1 | NC_056660.1 | 80.064 | 938 | 167 | 15 | 39 | 971 | 897429 | 898351 | 0 | 678 |
| OQ601562.1 | NC_056673.1 | 79.158 | 974 | 190 | 9 | 1 | 971 | 24734451 | 24735414 | 0 | 662 |
| OQ601562.1 | NC_056666.1 | 79.137 | 973 | 191 | 7 | 1 | 971 | 58091713 | 58092675 | 0 | 662 |
| Maza0008 | NC_010642.1 | 98.867 | 971 | 11 | 0 | 1 | 971 | 15188 | 16158 | 0 | 1733 |
| Maza0008 | NC_056676.1 | 90.628 | 971 | 91 | 0 | 1 | 971 | 5568675 | 5569645 | 0 | 1290 |
| Maza0008 | NC_056676.1 | 90.525 | 971 | 92 | 0 | 1 | 971 | 5585283 | 5586253 | 0 | 1284 |
| Maza0008 | NC_056666.1 | 79.725 | 947 | 184 | 4 | 1 | 947 | 58091713 | 58092651 | 0 | 678 |
| OQ629467.1 | NC_010642.1 | 98.866 | 970 | 11 | 0 | 1 | 970 | 15188 | 16157 | 0 | 1731 |
| OQ629467.1 | NC_056676.1 | 90.619 | 970 | 91 | 0 | 1 | 970 | 5568675 | 5569644 | 0 | 1288 |
| OQ629467.1 | NC_056676.1 | 90.515 | 970 | 92 | 0 | 1 | 970 | 5585283 | 5586252 | 0 | 1282 |
| OQ629467.1 | NC_056666.1 | 79.725 | 947 | 184 | 4 | 1 | 947 | 58091713 | 58092651 | 0 | 678 |
| OQ629468.1 | NC_010642.1 | 98.763 | 970 | 12 | 0 | 1 | 970 | 15188 | 16157 | 0 | 1725 |
| OQ629468.1 | NC_056676.1 | 90.619 | 970 | 91 | 0 | 1 | 970 | 5568675 | 5569644 | 0 | 1288 |
| OQ629468.1 | NC_056676.1 | 90.515 | 970 | 92 | 0 | 1 | 970 | 5585283 | 5586252 | 0 | 1282 |
| OQ629468.1 | NC_056666.1 | 79.725 | 947 | 184 | 4 | 1 | 947 | 58091713 | 58092651 | 0 | 678 |

We further examined the primers Wirdateti et al. (2024) used for PCR amplifications (forward: 5’-CTATAAGAACTTAATGACCAACATTCG-3’, reverse: 5’-TTCATTTAAGGAGGC GGTTTT-3’) using NCBI Primer-BLAST. The complete match of the primer pair is located in the tiger mtDNA (NC_010642.1). However, only 1 bp change in the forward primer sequence (5’-CTATAAGAAC **[T>C]** TAATGACCAACATTCG-3’) would make a complete match to a region on tiger chromosome F2 (NC_056676.1:5,568,577-5,569,727). This region encompasses the sequence that is the best BLAST match to OQ601561.1 and OQ601562.1 (NC_056676.1:5,568,675-5,569,645), the source of OQ601561.1 and OQ601562.1 from Numt amplifications.

## 3. Few details concerning quality control are provided to exclude the possibility of cross-contamination

The Numt problem alone which is systematic across the two specimens does not necessarily affect the authors’ conclusion regarding the presence of the Javan tiger. If the data are real, not cross-contamination, it is still possible that the hair collected belongs to the same group as the museum Javan tiger specimen. However, we are not able to find in the article if and how the authors excluded the possibility of contamination. What if the sequence was derived from contamination by the other control specimens? Were the DNA extraction and downstream experiments handled with the extreme precaution that is required for working with degraded genetic materials? Was the experiment replicated? Too few details regarding the quality control were provided to rule out the possibility of contamination during the experimental process.

The enormously high variant rate in the data prompted a concern that the sequence could be from contamination, not the single hair sample collected on site. There are 24 mismatches, out of the 971 bp sequence in length, between the hair and the museum Javan tiger specimen, corresponding to a genetic distance of 2.473×10^-2^. In the population genomic analyses including all tiger subspecies, only 196 variants were found across the mtDNA (15.5 kb in length with the control region removed) which is about 12.6 variants per 1,000 bp (Liu et al., 2018). The genetic difference among the Javan, Bali, and Sumatran tigers from the Sundaland, is even less with 44 variants across the 15.5 kb mtDNA sequence, corresponding to about 2.84 variants per 1,000 bp (Sun et al., 2023). For the tiger nuclear DNA, the single nucleotide variant (SNV) rate in different subspecies varies, ranging between 0.026% and 0.072%, which is 0.26-0.72 variants per 1,000 bp (Liu et al., 2018). Regardless of their origin from mitochondrial or nuclear DNA, the presence of such a large number of variant sites found between the “Javan tiger” sequences generated by the authors is unusual for two homologous sequences that are both from tigers and strongly indicative of data unreliability.

The errors may result from various reasons that are impossible to trace based on the information provided by Wirdateti et al. (2024). Nevertheless, considering the likelihood of contamination during the production of OQ601561.1 and OQ601562.1, it is inappropriate to use these sequences to conclude the existence of the Javan tiger.

Lastly, if the authors had provided detailed images of the hair, a morphological examination could have been performed.

The claim of the rediscovery of the Javan tiger by Wirdateti et al. (2024) has garnered widespread attention in the general public, as well as among scientists and conservationists. All of us would be thrilled to learn that the Javan tiger survives. We agree with the authors that “Whether the Javan tiger still occurs in the wild needs to be confirmed with further genetic and field studies.” Regrettably, the authors’ initial conclusions based on DNA analysis of one putative tiger hair sample are more likely to have arisen erroneously due to flawed experimental design and lack of scientific stringency than from an extant tiger. Clear and reliable visual, physical, or solid genetic evidence will be required to suggest that the Javan tiger still survives on the Island of Java nearly half a century since the last positive confirmed sighting.

## References

Jackson, P. & Nowell, K. (2008) Panthera tigris ssp. balica: The IUCN Red List of Threatened Species 2008: e.T41682A10510320. 10.2305/IUCN.UK.2008.RLTS.T41682A10510320.en [accessed 2 April 2024].

Kim, J.-H., Antunes, A., Luo, S.-J., Menninger, J., Nash, W.G., O’Brien, S.J. & Johnson, W.E. (2006) Evolutionary analysis of a large mtDNA translocation (numt) into the nuclear genome of the Panthera genus species. Gene, 366, 292–302.

Liu, Y.-C., Sun, X., Driscoll, C., Miquelle, D.G., Xu, X., Martelli, P., et al. (2018) Genome-wide evolutionary analysis of natural history and adaptation in the world’s tigers. Current Biology, 28, 3840–3849.

Luo, S.-J., Kim, J.-H., Johnson, W.E., Walt, J. Van Der, Martenson, J., Yuhki, N., et al. (2004) Phylogeography and genetic ancestry of tigers (Panthera tigris). PLoS Biology, 2, e442.

Seidensticker, J. (1987) Bearing witness: observations on the extinction of Panthera tigris balica and P. t. sondaica. In Tigers of the World: The Biology, Biopolitics, Management, and Conservation of an Endangered Species (eds R.L. Tilson & U.S. Seal), pp. 1–8. Noyes Publications, Park Ridge, NJ.

Sun, X., Liu, Y.-C., Tiunov, M.P., Gimranov, D.O., Zhuang, Y., Han, Y., et al. (2023) Ancient DNA reveals genetic admixture in China during tiger evolution. Nature Ecology & Evolution, 7, 1914–1929.

Wirdateti, W., Yulianto, Y., Raksasewu, K. & Adriyanto, B. (2024) Is the Javan tiger Panthera tigris sondaica extant? DNA analysis of a recent hair sample. Oryx, 1–6.

Xue, H.-R., Yamaguchi, N., Driscoll, C.A., Han, Y., Bar-Gal, G.K., Zhuang, Y., et al. (2015) Genetic ancestry of the extinct Javan and Bali tigers. Journal of Heredity, 106, 247–257.

